# *Plasmodium falciparum* derived extracellular vesicles reprogram host MAP Kinase signalling cascade to promote cytoadherence

**DOI:** 10.64898/2026.05.12.724568

**Authors:** Madhavi Sanjay Shukla, Reetesh Raj Akhouri, Suchi Goel

**Affiliations:** Indian Institute of Science Education and Research (IISER) Tirupati, India; Okinawa Institute of Science and Technology and Graduate University, Japan; GITAM University, Hyderabad, India

**Keywords:** Malaria, Parasite derived Extracellular vesicles, Cytoadherence, MAPK, miRNAs

## Abstract

Severe malaria pathogenesis, caused by *Plasmodium falciparum*, is linked to cytoadherence, during which parasitized RBCs bind to host endothelial receptors, causing microvascular obstruction. Extracellular vesicles released by *P.falciparum*-infected RBCs (pEVs) are found in higher numbers in the plasma of severe malaria patients, yet their role in cytoadherence remains unexplored. We investigated how pEVs from virulent (FCR3) and non-virulent (3D7A) parasite strains modulate host cell signalling cascade to guide parasite-host cell interactions. Here, we show that FCR3-EVs enhance the cytoadherence of both FCR3 and 3D7A by activating the MAP kinase pathway through p-ERK and downstream transcription factors, c-fos and c-jun. This resulted in upregulation of ICAM-1 and CD36, which promoted parasite binding with host cells. Moreover, pharmacological blockade of host cellular signalling responses abrogated cytoadherence, thereby defining a clear relationship between host cell signalling responses and cytoadhesion. Further, this modulation is due to human derived miRNAs in the pEVs that target phosphatase PTPRR; a negative regulator of MAP kinase leading to accumulation of pERK. However, 3D7A-EVs differ from FCR3-EVs as they neither activated c-fos and c-jun nor enriched miRNAs against PTPRR in cells. Therefore, our study provides new insights into the molecular mechanism underlying EV mediated parasite-host communication.

## Introduction

Malaria remains a significant health problem with an estimated 600,000 deaths annually in the tropical and subtropical parts of the world [1]. Among all *Plasmodium* species, *Plasmodiumfalciparum*is clinically the most devastating parasite due to its severe malaria manifestations that arise because of its ability to sequester in the microvasculature [2,3]. The most lethal hallmarks of severe malaria are cerebral malaria, severe anemia, respiratory distress, metabolic acidosis, and multiple organ failure. These symptoms result from the blockage of blood flow in organs either due to cytoadherence of parasitized RBCs (pRBCs) mediated by binding of PfEMP1s with ICAM-1, CD36, PECAM, VCAM-1, EPCR on endothelial cell surface or by rosetting where PfEMP1 binds with heparan sulfate, CR1, blood group antigens, and glycophorin B present on erythrocytes [2,4]. The most studied host receptor ICAM-1 is known to play a role in cerebral malaria [5], while EPCR is linked with respiratory distress symptoms [6] and gc1qR/HAPB1/p32 with seizures [7]. Among all the receptors, most of the *P. falciparum* clinical isolates depend either on ICAM-1 or CD36 for cytoadhesion [5]. However, clinical isolates from children could use multiple receptors for binding which not only compounds the problem but also suggests that a receptor cannot be linked to one hallmark of severe malaria. Thus, it is speculated that parasites either express a repertoire of PfEMP1s on their surface or a PfEMP1 can bind with different receptors using its various DBL or CIDR domains. Besides, the pathogenesis is also aggravated as ICAM-1 has been shown to increase on brain endothelial cell surface during cerebral malaria [8,9]. Overall, the cytoadhesion is a complex mechanism and it is important to understand how parasites drive this severity due to multiple receptor binding.

Moreover, it is not clear how VAR2CSA expressing parasites alter signalling cascade in endothelial cell line, HBEC-5i to induce the expression of cytokines, nucleosome assembly genes and anti-viral immune response molecules [10]. In conjunction with this, parasites possess PfPTP2 that is required to produce parasite derived extracellular vesicles (pEVs) [11] and upon uptake in endothelial cells enriches miR-451a that target ATF-1 and CAV-1 and modulate endothelial cell barrier properties [12]. This suggests that probably cytoadhering parasites can communicate and influence the host physiology through pEVs.

pEVs are approximately 150-300 nm size vesicles containing approximately 153 proteins from *P.falciparum*[13–14]. Proteome analysis detected virulence-associated proteins, heat shock 70 kDa protein, Pf-enolase, and Pf-actin at higher levels. They also encapsulate 23 human RAB proteins that modulate intracellular membrane trafficking and cytokine secretion in immune cells [14]. Several small RNAs (Y-RNAs, vault RNAs, snoRNAs, piRNAs) and 120 plasmodial RNAs with regulatory functions make up for the nucleic acid content in EVs [15].

Since during severe malaria, pEVs are secreted 30-40% [14] higher compared to normal RBCs and they can impact host signalling, we hypothesized their role in modulating cytoadherence of the parasites. In our study, we show that EVs from a virulent parasite, FCR3, enhanced cytoadherence of parasites by upregulating the classical MAP kinase pathway through increased phosphorylation of ERK. This resulted in over-expression of ICAM-1 and CD36 on the cell surface, causing enhanced cytoadherence. Also, EVs from FCR3 increased virulence of non-cytoadherent 3D7A by transforming it into cytoadherent phenotype. We further show that pEVs specifically packaged miRNAs in EVs, and target inhibitors such as PTPRR, a phosphatase leading to the accumulation of p-ERK and activation of the pathway through decreased PTPRR.

## Materials and Methods

### Parasite culture in EV-free media

*Plasmodium falciparum* strains (FCR3 and 3D7A; obtained from MR4) were cultured in RPMI-1640 medium supplemented with O-positive human red blood cells (4% haematocrit), 30 μg/ml gentamycin, and 10% heat-inactivated O-positive human plasma. FCR3 was routinely selected on anti-IT4VAR60 antibodies. To eliminate contamination of EVs, human plasma was centrifuged at 4200 rpm for 10 min at 4 °C to remove cells. The supernatant was then collected, and ultracentrifugation was performed at 100,000 × g for 2 h. This EV-depleted plasma was then used for culturing the parasite. The parasite were maintained with a gas mixture (90% nitrogen, 5% oxygen, 5% carbon dioxide) at 37 °C. Cultures were stained with acridine orange and visualized at 63X magnification under fluorescence microscope (Olympus) using the Alexa fluor 488 filter. The parasites were routinely synchronized using 5% sorbitol. The cultures were regularly tested for mycoplasma using the mycoplasma PCR kit.

### Mammalian Cell Culture

Human umbilical vein endothelial cells (HUVECs) and Chinese hamster ovary (CHO) cells were cultured at 37°C in a 5% CO_2_. CHO cells were maintained in RPMI media with 10% Fetal Bovine Serum (FBS) and 1% penicillin-streptomycin. Meanwhile, HUVECs were grown in endothelial cell growth media that consisted of Vascular Cell Basal Medium with VEGF-containing endothelial growth kits. The cultures were regularly tested for mycoplasma using the mycoplasma PCR kit.

### Binding assay with pRBCs

CHO or HUVEC cells were seeded at a confluency of 60-70% per well in 24-well plates. The spent media from FCR3, 3D7A at ∼15-20% parasitemia. For RBC-EVs, uninfected RBCs were incubated in parallel to parasite cultures. The spent media from parasite cultures, as well as uninfected RBCs, were collected and diluted at a ratio of 1:1 with the growth media of respective cells and added to the cells and incubated further for 24 h. For binding, the cells were washed 5X with incomplete media and parasite cultures at trophozoite stage (∼15% parasitemia) were added to each well and incubated for 1 h. The cells were washed 4X with incomplete media and the bound parasites were visualized using inverted microscope (Olympus) at 20X objective lens. For analysis, 10 fields were counted for each sample and the graphs were plotted as a box-whisker plot.

To understand the role of pEVs, cytochalasin D at a final concentration of 600 nM along with the spent media of pEVs, RBC-EVs as well as EV- was added to the cells and incubated for 24 h at 37℃at 5% CO_2_. After incubation, binding assays with the parasite cultures were performed. The bound parasites were visualized at 20X under inverted microscope. For each sample, 10 fields were counted and the graph was plotted as percent binding with respect to no inhibitor control.

Similarly, inhibitors for host signalling pathways, UO126 at final concentration of 5 μM and 10 μM, SB203580 at 1, 2.5 and 5 μM and SC75741 at 1, 2.5, 5 μM for CHO and 10 and 100 nM for HUVEC were used. The assays were performed as above.

For antibody inhibition assays, anti-PEMP1 antibodies at concentrations 250 μg/ml and 500 μg/ml and anti-ICAM-1 and anti-CD36 at 10 μg/ml were added along with the parasite cultures to the cells for 1 h at 37℃at 5% CO2. The cells were washed 4X with incomplete media and binding was visualized at 20X under inverted microscope (Olympus). For data analysis, 10 fields were counted per sample and the graph was plotted as percent binding with respect to no antibody control.

### Dynamic Light Scattering (DLS)

The hydrodynamic size distribution of isolated EVs was determined by dynamic light scattering. Following isolation and washing, freshly prepared pEVs from spent media of parasite cultures and RBC-EVs were gently resuspended in sterile, 0.22 µm filtered PBS by careful pipetting to ensure uniform dispersion and minimize vesicle aggregation. EV suspensions were subsequently serially diluted in filtered PBS to obtain the appropriate particle concentrations for analysis (50 μg/ml). Diluted samples were transferred into disposable polystyrene cuvettes and allowed to equilibrate at 25 °C before measurement. During acquisition, fluctuations in scattered light intensity resulting from the Brownian motion of the vesicles were recorded and used by the instrument software to calculate the hydrodynamic diameters. Particle size distributions were reported as intensity-weighted profiles, from which the mean hydrodynamic diameter and polydispersity index (PDI) were derived.

### Western blotting

Cells were lysed in 2× RIPA buffer, and the lysates were incubated on ice for 30 min. The lysates were centrifuged at 14,000 × g for 15 minutes and stored at –80 °C. The supernatants were then resolved by SDS–PAGE. Proteins were transferred to nitrocellulose membranes, which were blocked overnight at 4 °C. The membranes were washed three times with Tris-Buffered Saline + 0.5% Tween-20 and incubated with primary antibodies at a dilution of 1:1000 (p-ERK, ERK, IκBα, and β-tubulin) for 90 min. After 3X washing with TBST washes, the membranes were incubated either with anti-rabbit or anti-mouse HRP-conjugated at a 1:5000 dilution for 1 h at room temperature. The blots were washed 3X with TBST and developed with ECL substrate (Thermofisher) and visualized using chemiluminescence.

### Immunofluorescence assay

The coverslips were added to 6-well plates and cells at ∼60% confluency and used for immunofluorescence analysis after addition of RBC-EVs, pEVs. The coverslips were washed, fixed with 2% paraformaldehyde for 30 minutes, and incubated in blocking buffer (2% BSA in Phosphate-buffered saline, PBS). The fixed cells were then incubated with primary antibodies against the target proteins for 90 min, washed 5X with PBS, and incubated with either anti-mouse or anti-rabbit Alexa Fluor 594 and DAPI for 1 h. After washing with PBS, coverslips were mounted on slides using antifade medium, and the edges were sealed with nail polish. The cells were visualized using Nikon microscope at 40X objective lens using appropriate filters.

### Quantitative RT–PCR

Total RNA was isolated using the Trizol method and chloroform/isopropanol extraction. To separate the phases effectively, TRIzol reagents were mixed with chloroform, vigorously shaken, and then incubated at room temperature. Centrifugation was then employed to separate the phases of the solution. Following the collection of the aqueous phase containing RNA, isopropanol was added to precipitate the RNA and washed with 75% ethanol, air-dried briefly, and resuspended in nuclease-free water. RNA was treated with DNase I to remove any genomic DNA, and RNA was purified RNA using the PureLink RNA Mini Kit spin column.

For cDNA synthesis, 500 ng of total RNA was incubated with random hexamers and dNTPs mix. The RNA was denatured at 65°C for 5 min and incubated on ice for 2-3 min. The RNA was reverse transcribed using the Superscript RT first strand synthesis kit, which was used at 50°C for 10 min, at 80°C for 10 min. The PCR mix for real-time RT-PCR was prepared by addition of 1 µl of cDNA, 200 µM gene-specific primers, 1X SYBR green mix (Thermofisher,USA) and nuclease-free water and the reaction was set up at 50°C for 2 min, 95°C for 2 min, 40 X 95°C for 15 sec, 60°C for 1 min. The expression was normalized using GAPDH as the housekeeping gene and the fold change was calculated with respect to EV-using the 2^-ΔΔCt^ method.

### miRNA seq of pEVs and RBC-EVs

For miRNA isolation, EVs were isolated using ultracentrifugation at 100,000 x g for 2 h from spent media from 600 ml of parasite cultures (FCR3 and 3D7A) and RBC-EVs. For miRNA isolation, TRIzol was added to EVs and purified for small RNAs through the PureLink miRNA Isolation Kit (Invitrogen). Eluted purified miRNAs were then quantified for miRNA seq.

miRNA seq was performed commercially. Briefly, miRNA libraries were prepared using NEBNext Multiplex Small RNA Library Prep Kit and final libraries were quantified with Qubit 4.0 fluorometer using DNA HS assay kit. For identifying the insert size of the library, Tapestation 4150 was used, utilizing sensitive D1000 screentapes, following manufacturer’s protocol. Once the insert size ranged around ∼60-70, the library was sequenced using illumina and the data was analyzed. After obtaining the differential data, we considered log_2_CPM and analyzed targets for KRAS, MAPK1, DUSP1, DUSP4 and PTPRR using mirDB, TargetSCAN and DIANA tools. The miRNAs which were common in all the three softwares were taken for realtime RT-PCR analysis to check for their enrichment in cells.

### miRNA analysis

miRNA was isolated using above method from CHO cells incubated with RBC-EVs, FCR3-EVs, 3D7A-EVs along with EV- as control. The miRNA was reverse transcribed to cDNA using the MIRRT MystiCq microRNA cDNA Synthesis Kit, where miRNAs were polyadenylated and converted to cDNA using an oligo dT primer with a universal sequencing adaptor that allows universal primer amplification together with miRNA-specific primer sequences. Real-time quantitative PCR analysis was performed using SYBR Green dye with a primer concentration of 20 μM. The miRNA expressions were quantified as a ratio of normalised values SNORD44 and fold change was calculated with respect to EV- using the 2^-ΔΔCt^ method.

### Fluorescence-activated cell sorting

After incubation with RBC-EVs, pEVs and EV- as a control with CHO, the cells were scrapped using a cell scrapper and used for FACS analysis. Unstained cells and only anti-mouse Alexa Fluor 594 was used as negative controls for gating in FACS. The cells were washed 3X with PBS and incubated with anti-ICAM-1 and anti-CD36 antibodies at a concentration of 50 µg/mL in 2% BSA for 1 h at room temperature. The cells were washed with 4X with PBS and incubated with anti-mouse Alexa fluor-594 at a dilution of 1:100 for 1 h at room temperature. The cells were washed 5X with PBS and the data was collected with FACS-Celesta and 10,000 events were recorded for each condition. For data accquisition, unstained cell were used to set up the gate for cells while anti-mouse Alexa Fluor 594 was used to set up gate for anti-ICAM-1 and anti-CD36 staining. The data was analyzed using BD software and histograms were plotted R software.

For FACS analysis of the parasites, FCR3 and 3D7A cultures at the trophozoite stage at ∼15% parasitemia was washed 3X with PBS and staied anti-IT4VAR60 and anti-CIDRγ2 antibodies at a dilution of 50 µg/mL for 1 h. The parasites were washed again with PBS and incubated with anti-rabbit Alexa Fluor 488 and ethidium bromide at a concentration of 2.5 µg/mL for 1 h at room temperature. After washing 5X with PBS, the parasites were resuspended in PBS and the data accquisition was done in FACS-Celesta. For gating of RBCs, unstained parasite cultures were used. The parasites stained with only ethidium bromide was used to set up gate for parasites and for staining with antibody, 10,000 parasites were counted. As a negative control, anti-rabbit Alexa fluor 488 was used. The data was analyzed using FACS-celesta software.

## Results

### Spent media from parasite cultures increase the binding of FCR3 and 3D7A with host cells

To decode the role of pEVs in enhancing cytoadherence, we initially tested the binding of FCR3 and 3D7A parasites with Chinese Hamster Ovary (CHO) cells, which possesses multiple receptors for cytoadherence and the Human Umbilical Vein Endothelial cells (HUVEC) as an endothelial cell line (Figure 1A). In both CHO and HUVEC, FCR3 pRBCs consistently bound in large numbers, while 3D7A bound in low numbers with CHO and did not bind with HUVEC (Figure 1). Based on the avid binding of FCR3, we classified it as virulent and 3D7A as a non-virulent parasite line for cytoadherence.

**Figure 1.**
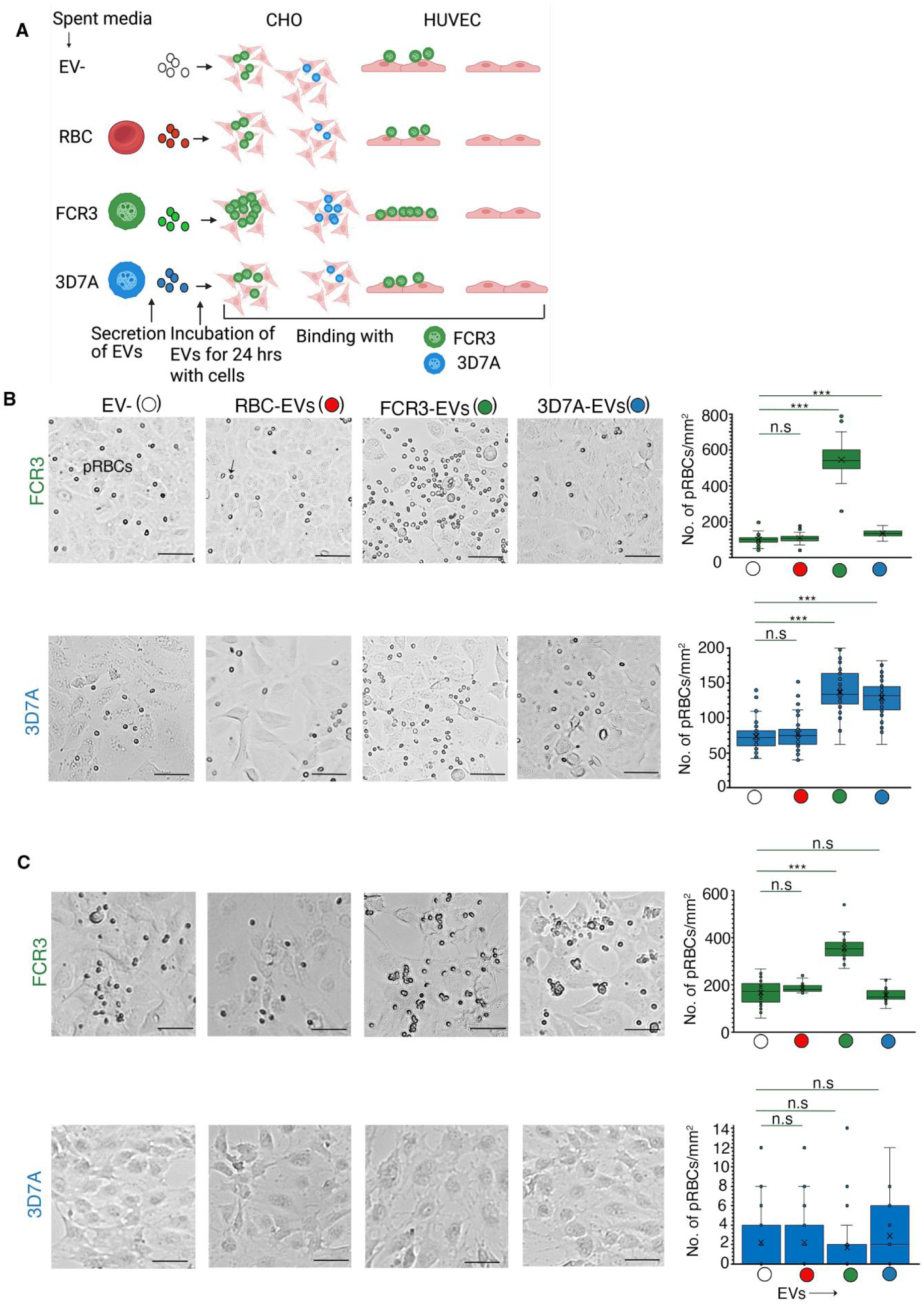
*P. falciparum* spent media modulate strain-specific cytoadherence of *P. falciparum*–infected erythrocytes. **(A)** Schematics depicting the experimental workflow to understand the role of EVs from virulent (FCR3) and non-virulent (3D7A) parasites in enhancing the cytoadherence. **(B)** and **(C)** shows representative images of FCR3 and 3D7A binding with CHO and HUVEC respectively along with box whisker plots of binding of FCR3 (green) and 3D7A (blue) upon incubation with FCR3-EVs (green), 3D7A-EVs (blue). RBC-EVs (red) and no EVs (EV-, white) were used as controls. The images were captured in Olympus microscope with 20X objective lens and the bound pRBCs were counted in 10 fields per experiment. With n=6, each point represents the bound parasites in a field/mm^2^ and the data is plotted as box whisker plot and line represents the median. Scale bar = 50 nm. Statistical significance was calculated using One Way ANOVA. *** represents p<0.001, n.s - p>0.05.

Since EVs are secreted in the growth media, thus in order to assess their role, we cultured the parasites till 32-34 hours post-invasion in EV-free complete medium and added to the host cells. Also, to further establish that the effect is specific to the spent media of parasites, we incubated uninfected RBCs in EV free media (RBC-EVs) in parallel and used them as control, along with EV free media (EV-, Figure 1A). When compared with EV- in CHO cells, FCR3-EVs increased FCR3 pRBCs binding by 5-fold, and 3D7A binding by 2-fold (Figure 1B). In contrast, the addition of 3D7A-EVs did not influence FCR3 binding but did augment 3D7A binding by 2-fold. However, RBC-EVs did not influence both FCR3 and 3D7A binding (Figure 1B).

In HUVEC, addition of FCR3-EVs enhanced the cytoadherence of FCR3 pRBCs by 2-fold however, 3D7A still did not show binding. With 3D7A-EVs, we observed no changes in FCR3 binding with HUVEC (Figure 1C).

### pEVs present in spent media are responsible for enhanced cytoadherence

To confirm that the observed effects were indeed due to pEVs and not due to other media components, we isolated EVs by ultracentrifugation and resuspended them in equivalent volumes of EV-free medium (Figure 2A). When we added the resuspended fraction of pEVs to cultured CHO and HUVEC cells, the binding increased by the same fold as the spent media (Figure 2A). In contrast, EV-free medium alone did not alter parasite binding (Figure 2A).

**Figure 2.**
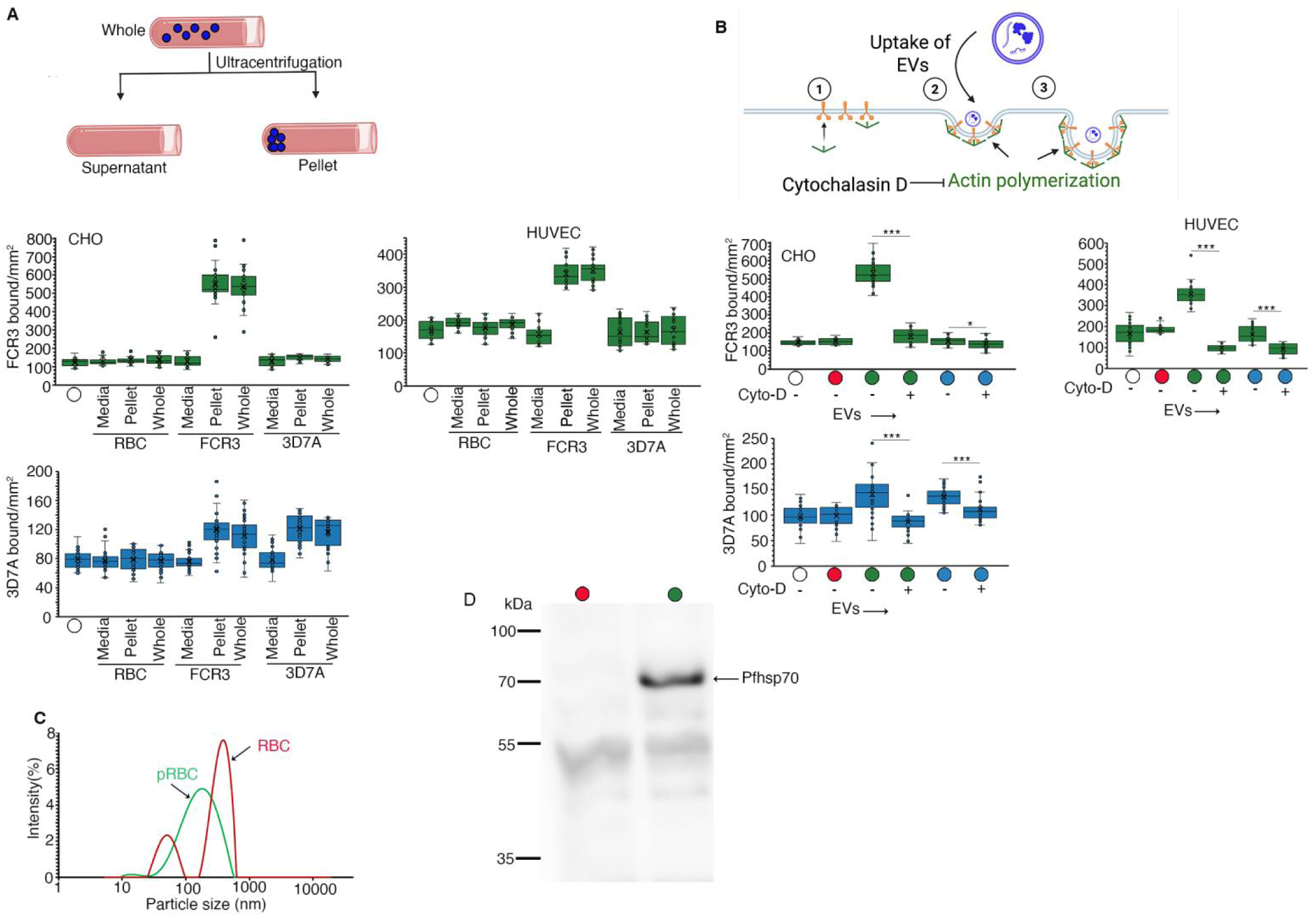
pEVs secreted in the spent media are required for enhanced cytoadherence. **(A)** Schematics (top) along with the box-and-whisker plots (bottom) of the bound FCR3 (green) and 3D7A (blue) to CHO and HUVEC under conditions; spent medium devoid of EVs (supernatant) and resuspended EVs (pellet). The spent media without ultracentrifugation (whole) is used as a positive control and EV- as a negative control. The images were captured in the Olympus microscope with 20X objective lens. With n=3, the bound pRBCs were counted in 10 fields per experiment and depicted as bound pRBCs/mm^2^ and the data is plotted as box whisker plot where line represents the median. **(B)** Illustration as shown in the top depicts that cytochalasin D blocks the actin polymerization that is required for active uptake of EVs in the cells. The bottom panel shows the binding of FCR3 (green) and 3D7A (blue) parasites on CHO and HUVEC cells with and without cytochalasin D upon incubation with FCR3-EVs (green), 3D7A-EVs (blue). RBC-EVs (red) and EV- (white) represent basal binding. n=3, the data is plotted as a box-whisker plot with each point representing bound pRBCs in a single field/mm^2^. Statistical significance was calculated using One way ANOVA. * represent p<0.05, *** represents p<0.001 **(C)** Dynamic light scattering profile displaying the particle-size distribution curves for EVs derived from RBCs and FCR3. The graph is a representation of two independent experiments (n=2) **(D)** Western blot of RBC-EVs and pEVs using antibodies against the parasite EV marker protein PfHSP70. The blot is a representation of three independent experiments (n=3).

Since EV uptake is an active, actin-dependent process, when we added cytochalasin D (an inhibitor of actin polymerization) along with EVs, the increased binding observed with FCR3- and 3D7A-EVs was abolished in both cell types, suggesting that active uptake of pEVs in host cells is essential (Figure 2B). Together, they confirmed that pEVs are solely responsible for the increased cytoadherence of pRBCs.

To characterize these vesicles, we performed dynamic light scattering (DLS) that measured an average vesicle size of ∼170 nm for pEVs and ∼ 250 nm for RBC-EVs (Figure 2C), consistent with previous reports on exosomes [14]. To confirm that these vesicles have originated from the parasite, we resolved the contents of vesicles on SDS-PAGE and performed western blot analysis using parasite-specific anti-PfHsp70 antibodies, a marker of parasite-derived exosomes. In the western blot, pEVs stained positive for Pf-Hsp70, with no such detection in RBC-EVs, confirming that most vesicles in the spent media were parasite-derived (Figure 2D). Based on their size and marker profile, we concluded that these vesicles are parasite-derived exosomes.

### Enhanced pRBC binding is mediated by PfEMP1

PfEMP1s primarily mediate parasite binding with host receptors, therefore we investigated their involvement using specific antibodies (Figure 3A). Based on our previous studies, where we have shown that FCR3 expresses IT4VAR60 and 3D7A expresses Pf3D7_0412900 [4], we used anti-IT4VAR60 and anti-CIDRγ2 of Pf3D7_0412900 to analyze the role of PfEMP1s in enhanced cytoadherence. We first examined whether these antibodies blocked basal parasite binding with CHO and HUVEC in EV- and RBC-EVs. The antibodies showed no inhibition of FCR3 and 3D7A binding with CHO and HUVEC cells at concentrations of 250 and 500 μg/ml, respectively (Figure 3B). Upon incubation with FCR3-EVs, the binding of FCR3 with CHO was blocked respectively, by 80% and 50% with anti-IT4VAR60 and anti-CIDRγ2 antibodies. However, neither of the antibodies blocked FCR3 binding upon incubation with 3D7A-EVs nor 3D7A binding after the addition of FCR3 and 3D7A-EVs (Figure 3B).

**Figure 3.**
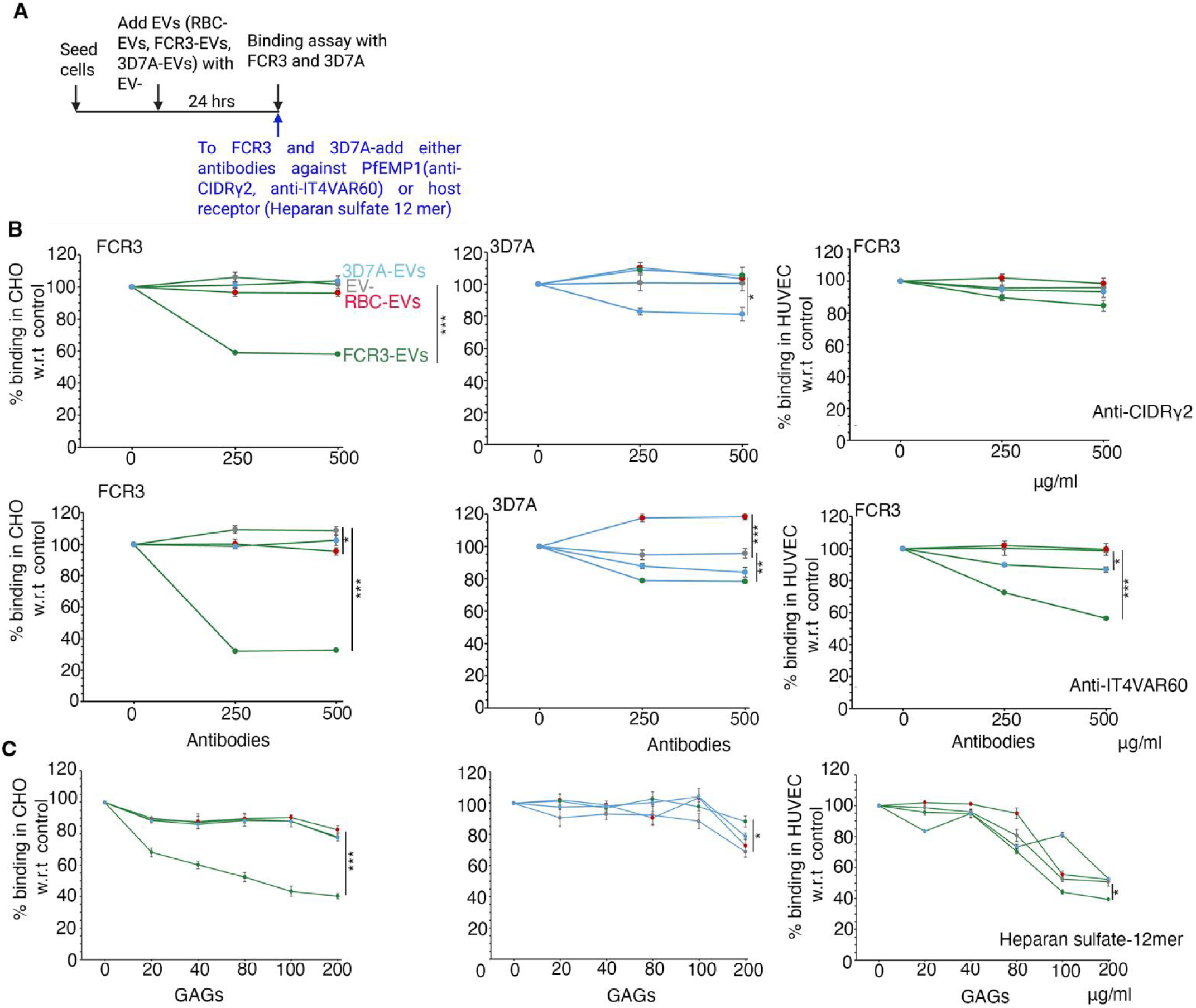
PfEMP1 dependent binding of malaria parasites after incubation with pEVs. **(A)** Schematics of the workflow of the experiment. **(B)** Antibody inhibition assays using anti-CIDRγ2 of PFD3D7_0412900 and anti-IT4VAR60 antibodies at concentrations of 250 μg/ml and 500 μg/ml. The plot shows the binding of FCR3 (green) and 3D7A (blue) with CHO (left) and HUVEC (right) upon addition of FCR3-EVs (green), 3D7A-EVs (blue) with RBC-EVs (red) and EV- (grey) as control at different concentrations of anti-CIDRγ2 of PF3D7_0412900 (top) and anti-IT4VAR60 (bottom) antibodies. With n=3, 10 fields per experiment were counted and the data is represented as line plots with mean percentage binding with respect to no antibody control. Data is presented as mean ± S.E.M and statistical significance was calculated using One way ANOVA.* represent p<0.05, ** represents p<0.01 and *** represents p<0.001, wherever not mentioned, the value of p>0.05. **(C)** Competition assay using receptor analog heparan sulfate-12mer. The plot depicts the binding of FCR3 (green) and 3D7A (blue) with CHO (left) and HUVEC (right) upon addition of FCR3-EVs (green), 3D7A-EVs (blue) with RBC-EVs (red) and EV^-^ (grey) as control after addition of 20, 40,80,100 and 200 μg/ml of heparan sulfate 12 mer. With n=3, 10 fields per experiment were counted and the data is represented as line plots with mean percentage binding with respect to no receptor analog control. Data is presented as mean ± S.E.M and statistical significance was calculated using One way ANOVA.* represent p<0.05, *** represents p < 0.001, wherever not mentioned, the value of p>0.05.

Further with HUVEC, FCR3 binding was reduced by anti-IT4VAR60 antibodies in a dose-dependent manner (∼45% and ∼40%, respectively) only in the presence of FCR3-EVs but not with 3D7A-EVs. However, unlike CHO cells, anti-IT4VAR60 did not completely block binding relative to the EV- control in HUVEC (Figure 3B).

Since anti-CIDRγ2 also blocked binding of FCR3, therefore we tested whether these antibodies recognize FCR3. FACS analysis showed that both anti-IT4VAR60 and anti-CIDRγ2 antibodies recognize FCR3 pRBCs but only anti-CIDRγ2 antibodies recognize 3D7A pRBCs (Figure S1).

Since IT4VAR60 uses heparan sulfate as the receptor [16], we also performed inhibition studies using soluble receptor analog, heparan sulfate 12 mer (Figure 3C). Similar to inhibition with anti-IT4VAR60, heparin sulfate 12-mer reduced binding of FCR3 with CHO by 50% only upon addition of FCR3-EVs with no effect on EV-, RBC-EVs and 3D7A-EVs (Figure 3C). However, this was not the case with HUVEC, where unlike anti-IT4VAR60 inhibition, heparin sulfate-12-mer independent of the origin of EVs showed an inhibition of 50% in FCR3 binding (Figure 3C). Thus, incomplete inhibition of binding indicated that there could be other receptors used by the parasites for cytoadherence.

### pEVs modulate signalling in host cells

Since the increased cytoadherence was specifically with pEVs that was partially inhibited with the receptor analog, thus we hypothesized that they might modulate host signalling cascade to upregulate receptors other than glycoaminoglycans. We focused on the MAPK/ERK and NFκβ pathways, which regulate expression of adhesins such as ICAM-1 and CD36 [17–20] linked to malarial parasite cytoadherence (Figure 4A). To track the activation of pathways, we performed western blot analysis for p-ERK and total ERK for the MAP kinase pathway, and IκBα for NFκβ, along with β-tubulin as a loading control. In all the cells, anti-ERK antibodies showed a double band pattern, probably for ERK1 and ERK2. Interestingly, we did not observe any changes in the level of ERK expression irrespective of treatment with pEVs, RBC-EVs or EV-. However, incubation with FCR3 and 3D7A-EVs increased the levels of p-ERK in CHO cells as compared with EV- but not with RBC-EVs (Figure 4B). To confirm whether the upregulation of p-ERK was due to uptake of pEVs, we tested its expression levels after the addition of cytochalasin D (Figure 4B). We found that cytochalasin D treatment reduced p-ERK levels, indicating the active involvement of pEVs in ERK signalling. Thus, no changes in the Iκβα levels suggest the role of the MAPK pathway and not the NFκB pathway. In HUVEC, pEVs did not alter Iκβα or p-ERK levels, where both ERK1 and ERK2 were expressed and phosphorylated at equal levels, suggesting no specific activation of the MAPK pathway with pEVs. In the absence of activation of ERK, we probed for the p38 pathway and found no changes upon addition of pEVs with respect to EV- or RBC-EVs (Figure S2). In contrast to previous studies where activation of the NFκβ pathway is suggested [10], we did not observe consistent down-regulation of Iκβα, an inhibitor of NFκβ, upon incubation of pEVs (Figure 4B).

**Figure 4.**
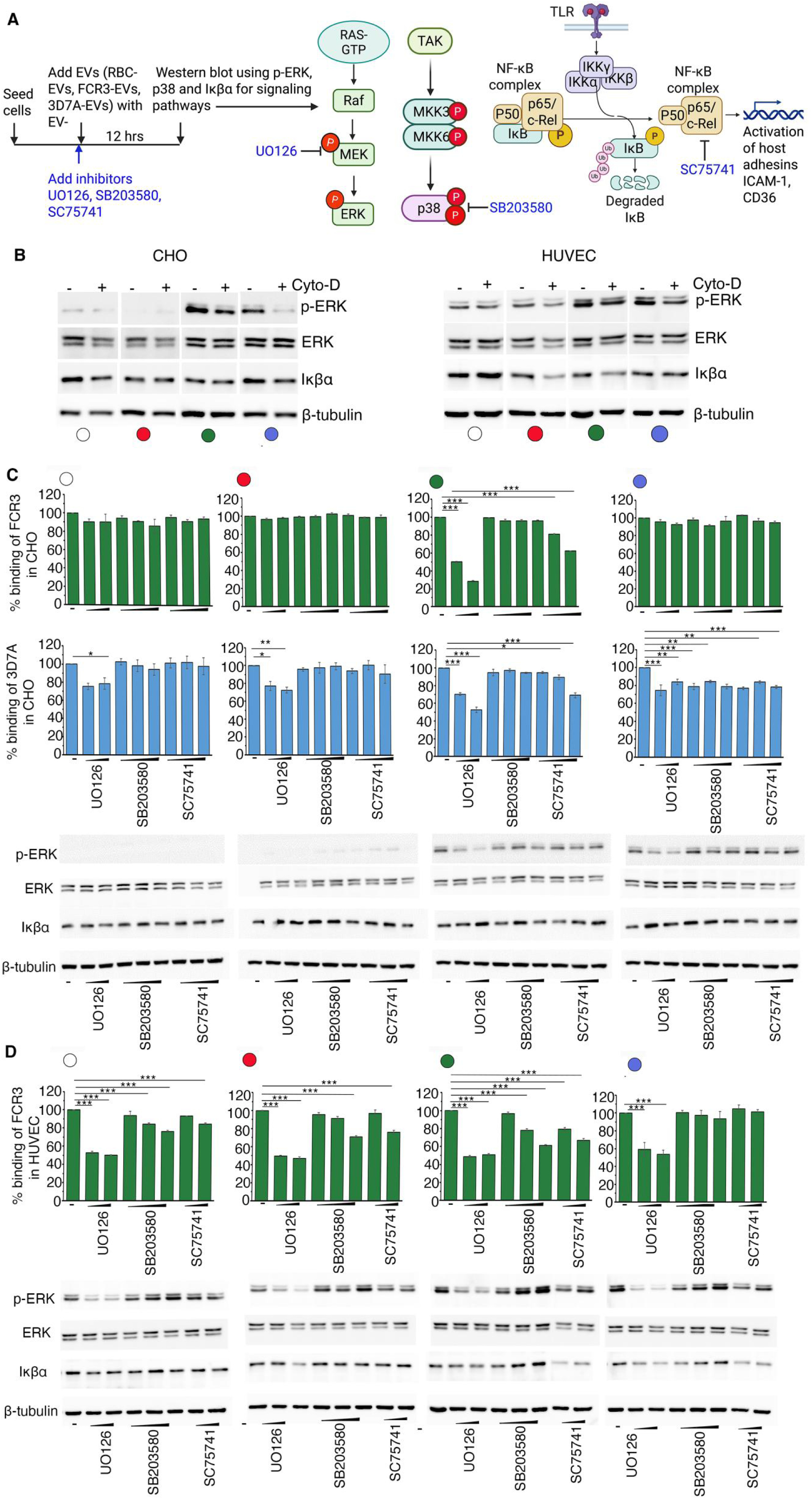
Activation of classical MAPK pathway upon incubation with pEVs. **(A)** Schematics depicting the workflow highlighting the MAPK, p38 and NFκβ pathway. The targets of the pharmacological inhibitors UO126, SB203580 and SC75741 for respectively MAPK, p38 and NFκβ are also marked in the schematics. **(B)** Immunoblot analysis in CHO and HUVEC cells with and without Cytochalsin D using anti-p-ERK, ERK, Iκβα after addition of RBC-EVs (red), FCR3-EVs (green), 3D7A-EVs (blue) along with EV- (white). β-tubulin was used as a loading control. The blot is a representation of four independent experiments (n=4). For **(C)** CHO and **(D)** HUVEC; The top panel shows the binding of FCR3 (green) and 3D7A (blue) upon addition of FCR3-EVs (green), 3D7A-EVs (blue) with RBC-EVs (red) and EV- (white) as control after addition of MEK inhibitor (U0126) at concentrations of 5 and 10 μM, p38 inhibitor (SB203580) at concentrations of 1, 2.5, 5 μM and NFκβ inhibitor (SC75741) at 1, 2.5, 5 μM. For HUVEC, 10 and 100 nM concentrations were used for SC75741. With n=3, 10 fields per experiment were counted and the mean percent binding in presence of inhibitors was calculated with respect to the no inhibitor control. The data is presented as mean ± S.E.M. Statistical significance was calculated using One way ANOVA. * represent p<0.05, ** represents p<0.01 *** represent p<0.001, wherever not mentioned, the value of p>0.05. The bottom panel is the immunoblot analysis of cells under the same conditions as the top panel, using antibodies against phosphorylated ERK (p-ERK), total ERK, IκBα, and β-tubulin as a loading control. The blot is a representation of three independent experiments.

To further investigate whether the activation of ERK signalling is directly responsible for increased parasite binding, we used specific inhibitors and tested their effects on parasite binding after adding pEVs (Figure 4A). We employed UO126, which inhibits MEK (upstream of ERK), SB203580 and SC75741 that target p38 MAPK and NFκβ pathway respectively (Figure 4A) [21]. Upon incubation of RBC-EVs as well as 3D7A-EVs with CHO, FCR3 binding did not reduce after addition of inhibitors which was similar to the effect of inhibitors on EV-. In contrast, 3D7A binding showed 10-20 % reduction in presence of inhibitors in EV-, RBC-EVs and 3D7A-EVs. Only when CHO cells were incubated with FCR3-EVs, UO126 at a 10 nM concentration brought FCR3 and 3D7A binding to levels similar to those of EV- (Figure 4C). SB203580 did not affect cytoadherence, whereas SC75741 caused ∼35% inhibition at the highest concentration (5 μM) in both the parasites (Figure 4C). With HUVEC, UO126 displayed an inhibition of ∼ 49% in FCR3 binding in EV- which was similar in RBC-EVs as well as pEVs. Moreover, incubation of FCR3-EVs with HUVEC also resulted in partial inhibition of ∼35% with SB203580 and SC75741 (Figure 4D).

Thus, the ability of UO126 to block FCR3 binding with CHO upon incubation of FCR3-EVs suggests the involvement of the classical MAP kinase pathway. However, in HUVEC, we did not observe any upregulation of p-ERK; these inhibitors probably blocked the basal signalling pathway and thus affected binding. To confirm this, we probed p-ERK and Iκβα levels by western blot analysis after adding inhibitors. Western blot analysis of CHO cells confirmed our binding assay results, where p-ERK levels were affected specifically by UO126. Further, UO126, SB203580, and SC75741 did not alter ERK or Iκβα levels compared to EV- (Figure 4C). When we probed similarly with HUVEC, UO126 inhibited p-ERK that was independent of the addition of EVs, while with SB203580 and SC75741, p-ERK, the total ERK and Iκβα remained unaltered (Figure 4D). These results align with the binding assay data, suggesting that downregulation of p-ERK reduces receptor levels as well as cytoadherence.

### MAPK pathway is differentially modulated upon addition of FCR3 and 3D7A-EVs

Although both FCR3-EVs and 3D7A-EVs activate p-ERK, but the distinct inhibition of binding with UO126 suggests that there exist differences in how they activate the MAP kinase pathway. Therefore, in order to understand the differences, we tracked the classical MAP kinase pathway, where we evaluated the expression of c-fos, a transcription factor which is the downstream effector of the classical MAP kinase pathway (Figure 5A) [22]. The incubation of FCR3-EVs led to a significant 4.5-fold increase in c-fos expression compared to EV- and RBC-EVs, specifically in CHO (Figure 5B). Although 3D7A-EVs phosphorylated ERK, c-fos expression remained at levels similar to those in controls (Figure 5B). In HUVEC consistent with the lack of p-ERK activation, FCR3-EVs and 3D7A-EVs did not induce c-fos expression (Figure 5B). Similarly using an immunofluorescence assay, we found that FCR3-EVs over-expressed c-fos, while 3D7A-EVs showed levels comparable to EV- and RBC-EVs (Figure 5C). Similarly, in HUVEC, no change in expression was observed with either FCR3-EVs or 3D7A-EVs, which was consistent with the real-time RT-PCR results (Figure 5C).

**Figure 5.**
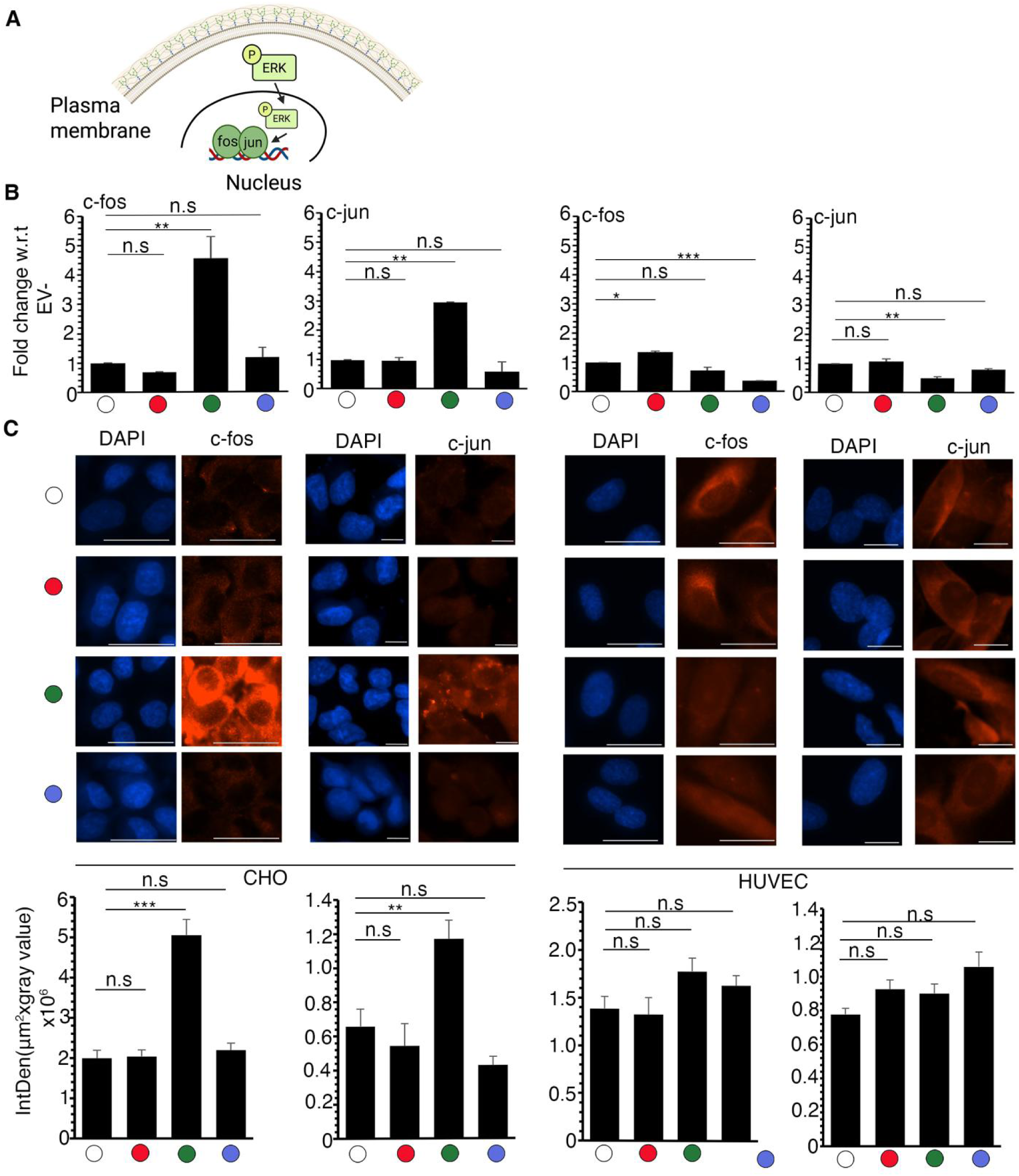
MAPK pathway is differentially modulated upon addition of FCR3 and 3D7A-EVs. **(A)** Illustration showing the downstream effectors of MAPK pathway where transcription factors c-fos and c-jun after activation binds to DNA to initiate transcription of host adhesins. **(B)** Realtime RT-PCR of c-fos and c-jun in CHO (left) and HUVEC (right) after addition of RBC-EVs (red), FCR3-EVs (green), 3D7A-EVs (blue) along with EV-(white). The expression was normalized with GAPDH and the fold change was calculated with respect to EV- using 2^-△△Ct^ method. With n=2 and performed in triplicates, the data is presented as mean ± S.E.M. Statistical significance was calculated using One-way ANOVA. * represent p<0.05, ** represent p<0.01, *** represent p<0.001, n.s represent p>0.05. **(C)** Immunofluorescence analysis of c-fos and c-jun (top) in CHO and HUVEC after addition of RBC-EVs (red), FCR3-EVs (green), 3D7A-EVs (blue) along with EV- (white). c-fos and c-jun was detected using Alexa fluor-594 and nuclei was stained using DAPI, n=3. Scale bars-For CHO c-fos, HUVEC c-jun- 25 µm and CHO c-jun CHO, HUVEC c-fos- 10 µm. Along with images, bar graph (bottom) depicts the integrated fluorescence intensity for c-fos and c-jun immunostaining in CHO and HUVEC cells. To quantify fluorescence, n=3, 5 images per experiment were captured in the microscope using a 40X objective lens using the same settings in fluorescence microscope (Nikon) with Texas Red filter. The image quantification was performed using ImageJ software and the data obtained is calculated as mean ± S.E.M. Statistical significance was calculated using One way ANOVA. ** represent p<0.01, *** represents p<0.001, n.s represents p>0.05.

Previous studies have shown that c-fos dimerizes with c-jun and binds to DNA [23], activating host adhesins transcription (Figure 5A). Therefore, we analyzed c-jun expression after adding pEVs. Similar to c-fos, real-time RT-PCR results indicate that addition of FCR3-EVs specifically increased c-jun transcription by three-fold in CHO cells compared to EV-, 3D7A-EVs and RBC-EVs (Figure 5B), which was also confirmed by immunofluorescence analysis (Figure 5C). In HUVEC, real-time RT-PCR and IFA results showed no change in c-jun levels with both FCR3- and 3D7A-EVs as compared to EV- and RBC-EVs (Figure 5B and C).

### Parasites use multiple host receptors for cytoadherence

Our experiments confirmed that FCR3-EVs activate transcription factors c-jun and c-fos that will finally lead to over-expression of host adhesins such as ICAM-1 and CD36 on the cell surface. In order to analyze this, we probed for the expression of these receptors using flow cytometry. We confirmed the increased surface expression of ICAM-1 only with FCR3-EVs, whereas CD36 expression was upregulated with both 3D7A-EVs and FCR3-EVs (Figure 6A), indicating the selective activation of receptor molecules by virulent parasite strains.

**Figure 6.**
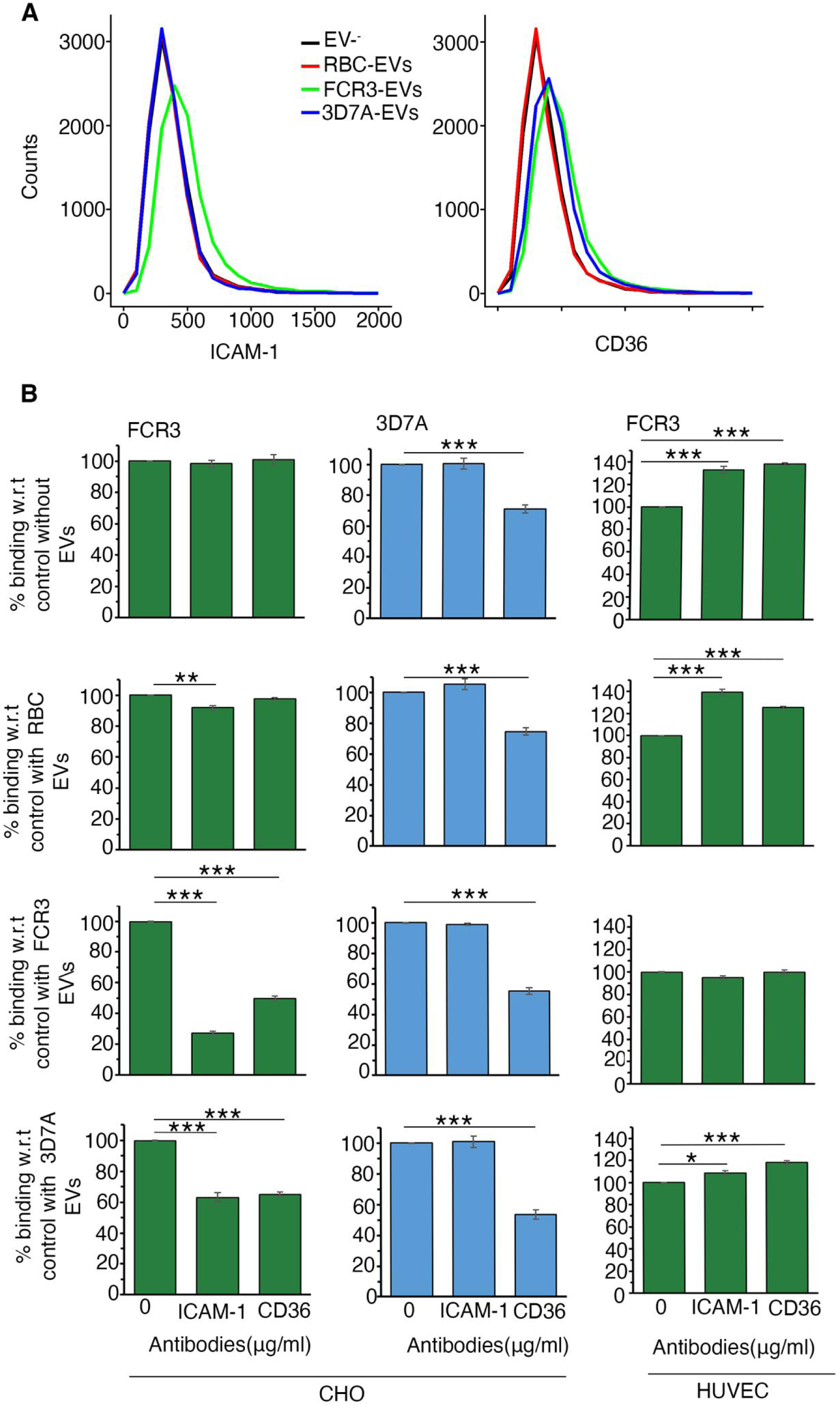
pEVs induce the expression of ICAM-1 and CD36 on CHO cell surface and is responsible for increased cytoadherence. **(A)** FACS analysis of ICAM-1 and CD36 on CHO cell surface after addition of RBC-EVs, FCR3-EVs and 3D7A-EVs with EV- as control. n=3, for each sample, 5000 cells were counted and plotted as a histogram. The histogram shown is a representation of three independent experiments. **(B)** The graphs depicting the percentage binding of FCR3 (green) and 3D7A (blue) obtained with anti-ICAM-1 and anti-CD36 antibodies at a concentration of 10 μg/ml each upon incubation with FCR3-EVs (green), 3D7A-EVs (blue) with RBC-EVs (red) and EV- (white) as control in CHO and HUVEC. With n=3, 10 fields per experiment were counted and the percentage binding was calculated with respect to the no antibody control. The data is calculated as mean ± S.E.M. Statistical significance was calculated using One Way ANOVA. * represent p<0.05, ** represent p<0.01, *** represent p<0.001, wherever not mentioned, p>0.05.

In order to test whether the upregulation of ICAM-1 and CD36 is responsible for enhanced cytoadherence, we performed inhibition with anti-ICAM-1 and anti-CD36 antibodies with both CHO and HUVEC. With anti-ICAM-1 antibodies, FCR3 binding with CHO was inhibited by 80% and 50% upon addition of FCR3-EVs and 3D7A-EVs respectively. When we tested the effect of anti-CD36 on FCR3 binding with CHO, it showed an inhibition of 50% with both FCR3- and 3D7A-EVs. Further no reduction of FCR3 binding with CHO in case of EV- and RBC-EVs suggests that the enhanced cytoadherence in FCR3 is mediated by the host receptors ICAM-1 and CD36. In contrast, 3D7A binding with CHO was reduced only with anti-CD36 antibodies that was independent of addition of EVs (Figure 6B). Further with HUVEC, no reduction of FCR3 binding was observed with both anti-ICAM1 and anti-CD36 antibodies that correlates with the non-activation of MAPK pathway (Figure 6B).

### miRNAs in pEVs regulate the expression of host adhesins

We show that pEVs upregulate the host MAP kinase pathway, leading to the overexpression of the receptors; ICAM-1 and CD36 resulting in enhanced cytoadherence. We initially probed if cytokines, IL-12 and TNF-α could induce receptor over-expression. Upon performing ELISA of supernatants collected after addition of pEVs, RBC-EVs and EV- to HUVEC, we did not detect any changes in levels of IL-12 and TNF-α when compared to EV-, suggesting that cytokines do not mediate this differential regulation of host adhesins (Figure S3). Therefore, we explored whether miRNAs of human origin present in pEVs could modulate MAP kinase pathways. To investigate this, we first checked whether activators or inhibitors of this pathway were upregulated or downregulated in the cells (Figure 7A). Using real-time RT-PCR, we analyzed the expression of activators; KRAS and MAPK [24,25] and inhibitors: DUSP1, DUSP4 (phosphatases specifically p38) [26,27], and PTPRR (protein tyrosine phosphatase) known to regulate activity of MAP kinases [28,29] (Figure 7A). Adding FCR3-EVs upregulated KRAS and MAPK in CHO cells as compared to EV-, although only at low levels (∼1.5-fold, Figure 7B). Further consistent with the non-activation of c-fos and c-jun with 3D7A-EVs, no activation of KRAS and MAPK1 was observed (Figure 7B). DUSP1 and DUSP4 did not change compared to EV- and RBC-EVs, however, FCR3-EVs and 3D7A-EVs especially downregulated PTPRR (Figure 7B). In HUVEC, we did not observe either upregulation of activators or downregulation of inhibitors consistent with the above results (Figure 7B). In order to establish whether miRNAs in EVs would play any role, we performed miRNA-seq of FCR3-EVs and 3D7A-EVs along with RBC-EVs as a control (Figure 7A). Upon differential analysis of the data, we observed both common and unique sets of miRNAs are enriched in FCR3-EVs and 3D7A-EVs which could interfere/target signalling pathways (Figure 7C, S4). Based on our results, here again we mostly focused on miRNAs that are distinctly present in FCR3-EVs and 3D7A-EVs and target the MAPK pathway (Figure 7D). Using miRDB, TargetSCAN and Diana to identify targets, from the list of miRNAs that are downregulated with respect to RBC-EVs, we could identify miRNAs that target KRAS and MAPK1. For FCR3-EVs, ∼ 8 miRNAs are downregulated for MAPK1/KRAS as compared to 3D7A-EVs where mir-548bc and mir-548z were downregulated. Similarly, from the list of miRNAs that are upregulated with respect to RBC-EVs, we could fish out miRNAs that target DUSP1, DUSP4 and PTPRR where both FCR3-EVs and 3D7A-EVs contain multiple miRNAs that target the three inhibitor genes (Figure 7D). Since after uptake of EVs, miRNAs should be released inside the cells, therefore we tested their enrichment in CHO using miRNA realtime RT-PCR. For miRNAs that target KRAS/MAPK1, we found that in FCR3-EVs multiple miRNAs; mir-6131, mir-4262, mir-1268a are significantly downregulated. While in 3D7A-EVs, mir-548bc was not detected and mir-548z did not show drastic reduction in the cells that is consistent with the expression levels of MAPK1 and KRAS in CHO (Figure 7E). Further based on the expression of inhibitors in CHO, we analyzed enrichment of PTPRR in CHO and observed that specifically in FCR3-EVs, the targets of PTPRR; mir-5010-5p and mir-6501-5p are upregulated (Figure 7E). Thus, the combination of downregulated PTPRR and over-expressed MAPK1 and KRAS leads to the activation of the p-ERK pathway.

**Figure 7.**
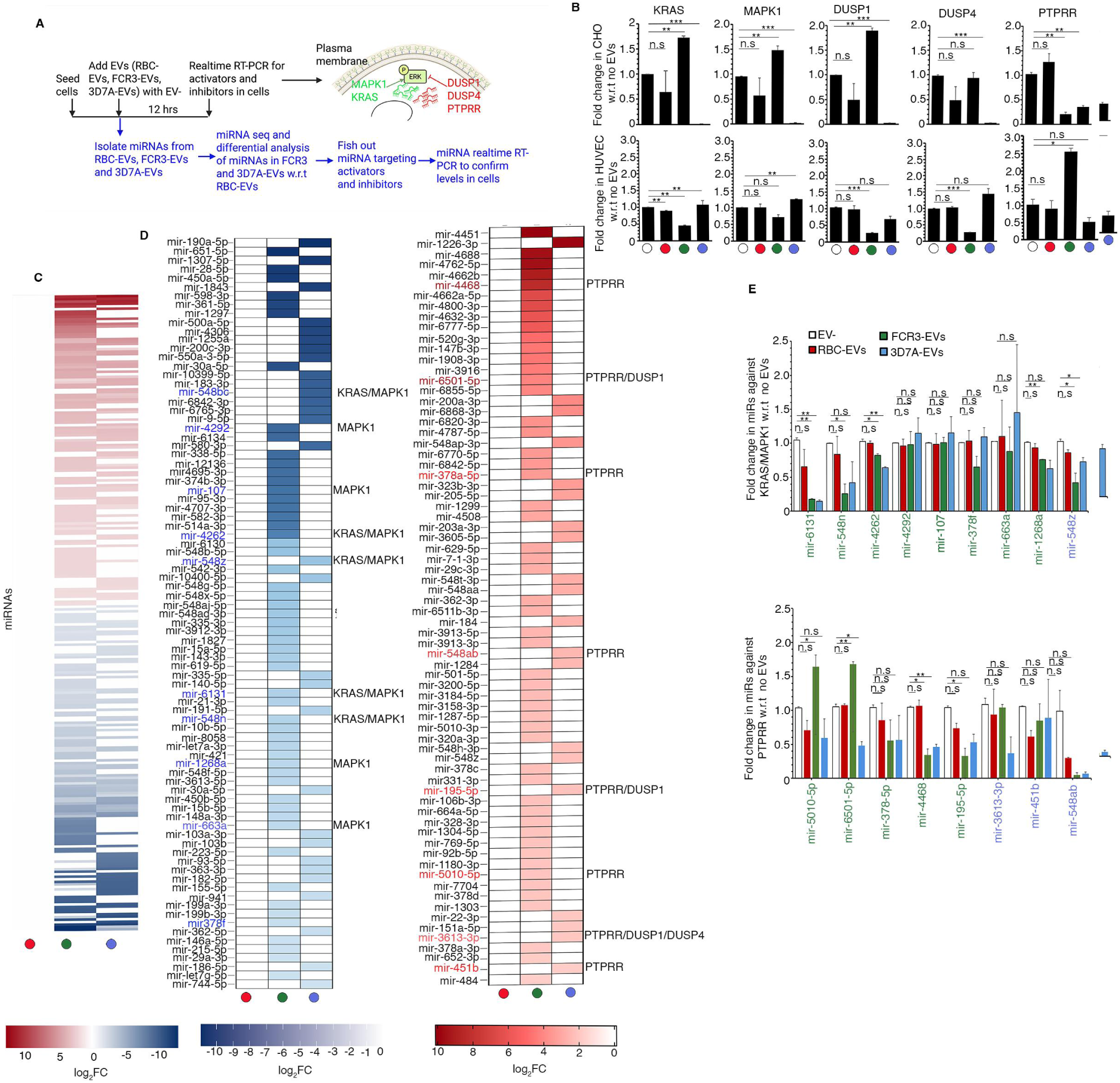
miRNAs modulate MAPK pathway. **(A)** Illustration depicting the workflow highlighting MAPK activators; KRAS and MAPK1 and inhibitors; DUSP1, DUSP4 and PTPRR **(B)** Real-time RT-PCR of KRAS, MAPK1, DUSP1, DUSP4 and PTPRR in CHO (top) and HUVEC (bottom) after incubation with RBC-EVs (red), FCR3-EVs (green) and 3D7A-EVs (blue) with EV- (white) as control. With n=2 and performed in triplicates, the expression was normalized with GAPDH and the fold change was calculated with respect to EV- using 2^-△△Ct^ method. The data is presented as mean ± S.E.M. Statistical significance was calculated using unpaired t-test. * represent p<0.05, ** represent p<0.01 *** represent p<0.001 and n.s-p>0.05. **(C)** Heat map of miRNAs represented as log_2_FC that are differentially present in FCR3-EVs (green) and 3D7A-EVs (blue) relative to RBC-EVs (red). **(D)** Heat map of unique miRNAs either downregulated or upregulated in FCR3-EVs (green) and 3D7A-EVs (blue) relative to RBC-EVs (red). miRNAs that target MAPK activators in downregulated miRNA heat map is colored as blue and miRNAs that target inhibitors in upregulated miRNA heat map are colored as red **(E)** Real-time RT-PCR of miRNAs that target KRAS/MAPK1(top) and PTPRR (bottom) after incubation of CHO cells with RBC-EVs (red), FCR3-EVs (green) and 3D7A-EVs (blue) and EV^-^ as a control. With n=2 and performed in triplicates, the expression was normalized with miRNA of SNODR44 and fold change was calculated with respect to EV- using 2^-△△Ct^ method. The data is presented as mean ± S.E.M. Statistical significance was calculated using unpaired t-test. * represent p<0.05, ** represent p<0.01, *** represent p<0.001 and n.s-p>0.05.

## Discussion

*P.falciparum*can modulate endothelial cell signalling pathways through either cytoadhesion that influences genes associated with NFκβ signalling, inflammation and angiogenesis [10] or parasite egress material capable of activating JAK STAT pathway [30]. Further, co-culture of parasites with cells was also shown to affect signalling in host cells [31], though responsible component could not be identified. Since pEVs are found in higher numbers in plasma of severe malaria patients [32], therefore we wanted to understand whether pEVs could impact signalling and enhance cytoadherence.

It has been established through miRNA and proteomics studies that pEVs specifically package parasite proteins and human derived miRNAs present in RBCs [33]; however, no reports suggest that components of EVs varies from one parasite strain to the other. In *Trichomonas vaginalis*, it was shown that EVs from virulent strain transformed non-virulent phenotype into virulent [34]. Thus, in our studies, we used EVs from virulent FCR3 and non-virulent 3D7A along with the RBC-EVs as a control. We demonstrate that EVs from spent media of FCR3 enhanced cytoadherence of FCR3 but importantly also of non-virulent 3D7A. As binding of parasites was not influenced with RBC-EVs and reversed with cytochalasin D, suggesting that the increased cytoadhesion is due to active uptake of pEVs by host cells and not by parasite egress material.

Further, the cytoadhesion of FCR3 was majorly sensitive to anti-IT4VAR60 and partially to anti-CIDRγ2 antibodies which is consistent with the surface recognition of FCR3 with both the antibodies. The recognition of FCR3 with anti-CIDRγ2 is probably due to the cross-reactivity with the CIDR domain in IT4VAR60. Though anti-CIDRγ2 from 3D7A showed surface reactivity but did not inhibit binding of 3D7A, suggesting that PF3D7_0412900 is not used for cytoadherence. In HUVEC, although partial inhibition was observed with anti-IT4VAR60 but no specific effect of heparan sulfate-12 mer was observed, suggesting that the increase obtained with FCR3-EVs is mediated by receptor other than heparan sulfate. Therefore, it is highly likely that pEVs upon uptake by host cells, modulate over-expression of host adhesins, such as ICAM-1 and CD36, on their surface. Upon tracking expression profile of signalling molecules in CHO and HUVEC, we observed specific phosphorylation of ERK upon uptake of EVs from FCR3 as well as 3D7A into CHO cells. This suggests that classical MAPK pathway is activated upon incubation with pEVs. However, pEVs did not bring any change in the levels of ERK phosphorylation in HUVEC. Alternatively, we also did not observe p38 activation. Further, Iκβα, inhibitor of NFκβ pathway showed no degradation, suggesting that relB is still sequestered with Iκβα and is not translocated to the nucleus for activation of the NFκβ pathway. Since in HUVEC, FCR3 binding is enhanced upon addition of FCR3-EVs, therefore we speculated whether cytokine secretion results into increased host adhesin expression and thus cytoadherence. However, no changes in the levels of TNF-α and IL-12 was observed with respect to EV-, raising the possibility that there could be other signalling factor(s) activated upon incubation with FCR3-EVs.

Although p-ERK is activated by both FCR3 and 3D7A-EVs, however only FCR3-EVs activated the transcription of c-fos and c-jun that dimerizes to initiate the transcription of host adhesins and overexpression of ICAM-1 and CD36 on the surface. This overexpression is responsible for enhanced cytoadherence as both anti-ICAM-1 and anti-CD36 antibodies reduced FCR3 binding to CHO cells.

3D7A-EVs, unlike FCR3-EVs, only promoted surface expression of CD36; probably activation of p-ERK can work with basal levels of transcription factors to cause this over-expression. This partial activation was responsible for cytoadherence as anti-CD36 antibodies brought down the FCR3 binding by 50%. In contrast to FCR3, 3D7A did not depend on ICAM-1 and CD36 for cytoadherence, suggesting they might require receptors other than ICAM-1 and CD36 for cytoadherence. In HUVEC, consistent with non-activation of transcription factors, no such inhibition with antibodies was observed. Together, these results also indicated that activation of signalling pathways is dependent on the strain that secretes pEVs as well as on the origin of cell types, which will finally guide which receptors are over-expressed on the cell surface.

Next, we wanted to understand how pEVs cause this modulation. We ruled out the role of cytokines, especially TNF-α, in ICAM-1 over-expression, as CHO did not secrete cytokines, while no specific secretion of IL-12 and TNF-α was observed upon addition of pEVs with respect to controls (EV-, RBC-EVs) in HUVEC. Sine miRNAs are packaged specifically in the case of pEVs [33], and their origin is human; therefore, we investigated whether miRNAs from pEVs modulate the host signaling cascade. Though it has been reported that certain miRNAs are enriched in pEVs, here we show that miRNAs differ between parasite strains.

Globally, miRNA population in pEVs targeted transcription factor binding but targets for the MAPK pathway were also present in both FCR3 and 3D7A. Based on the miRNA seq and experimental data, we focused on proteins that could regulate the pathway: activators (KRAS, MAPK1/ERK2) and inhibitors (DUSP1, DUSP4, PTPRR). KRAS functions as inducer of MEK that activates ERK for phosphorylation [35,36] while MAPK1 is a serine threonine kinase [37] DUSP1, DUSP4 [38] and PTPRR are phosphatases [39] and act as negative regulators of the MAPK pathway. Before identifying their targets in miRNA sequencing, we investigated their expression and found that KRAS and MAPK1 are upregulated upon addition of FCR3-EVs but not with 3D7A-EVs in CHO. In HUVEC, no changes were observed in KRAS and MAPK1 while DUSP1 and DUSP4 did not change in both the cell lines. However, we observed a significant reduction in PTPRR in CHO upon addition of FCR3- and 3D7A-EVs. Further when we analyzed whether miRNA targets are differentially expressed in pEVs, we found that KRAS and MAPK1 miRNAs are downregulated while PTPRR targets are upregulated in comparison to RBC-EVs. Interestingly, both FCR3-EVs and 3D7A-EVs contain targets of KRAS, MAPK1 and PTPRR, but miRNAs are different, suggesting that possibly this is the basic reason EVs have different capacity in terms of activation of signalling pathways in host cells. In order to confirm, we checked for the expression of miRNAs in CHO after addition of pEVs, where we found that mir-5010-5p and mir 6501-5p against PTPRR are specifically enriched in FCR3-EVs as well as CHO incubated with FCR3-EVs, but none of the miRNAs are enriched with 3D7A-EVs. Therefore, it is possible that although miRNAs are specifically present in FCR3 and 3D7A-EVs, the final activation of the MAP kinase pathway depends on which of the target miRNA are available for targeting.

The case with 3D7A-EVs is intriguing as p-ERK is phosphorylated, but c-fos and c-jun are present at the same levels as controls. Further KRAS and MAPK1 are clearly downregulated in 3D7A-EVs. On comparing the miRNA sequencing data, we found that 3D7A-EVs are also enriched with MAPK1 and KRAS miRNA, possibly the reason behind the downregulation of MAPK1 and KRAS. We speculate that ERK is phosphorylated through MEK1 and with basal levels of c-fos, is able to upregulate CD36 on the cell surface and thus is able to increase its own ability to cytoadhere.

## Conclusion

Overall, from our study we could establish that incubation of EVs from virulent strain FCR3 can modulate signalling pathways, leading to over-expression of host adhesins and thus promote the severity of malaria parasites through enhanced cytoadherence. Although 3D7A-EVs possess limited capability, but nonetheless it increases its own cytoadherence, clearly suggesting that pEVs can modulate cytoadherence and increase the virulence of the parasites. Overall, our studies provide a possible mechanism where EVs secreted by the malaria parasites lead to an increase in ICAM-1 levels and the multiple receptor binding phenotype observed in the clinical isolates of severe malaria patients (Figure 8).

**Figure 8.**
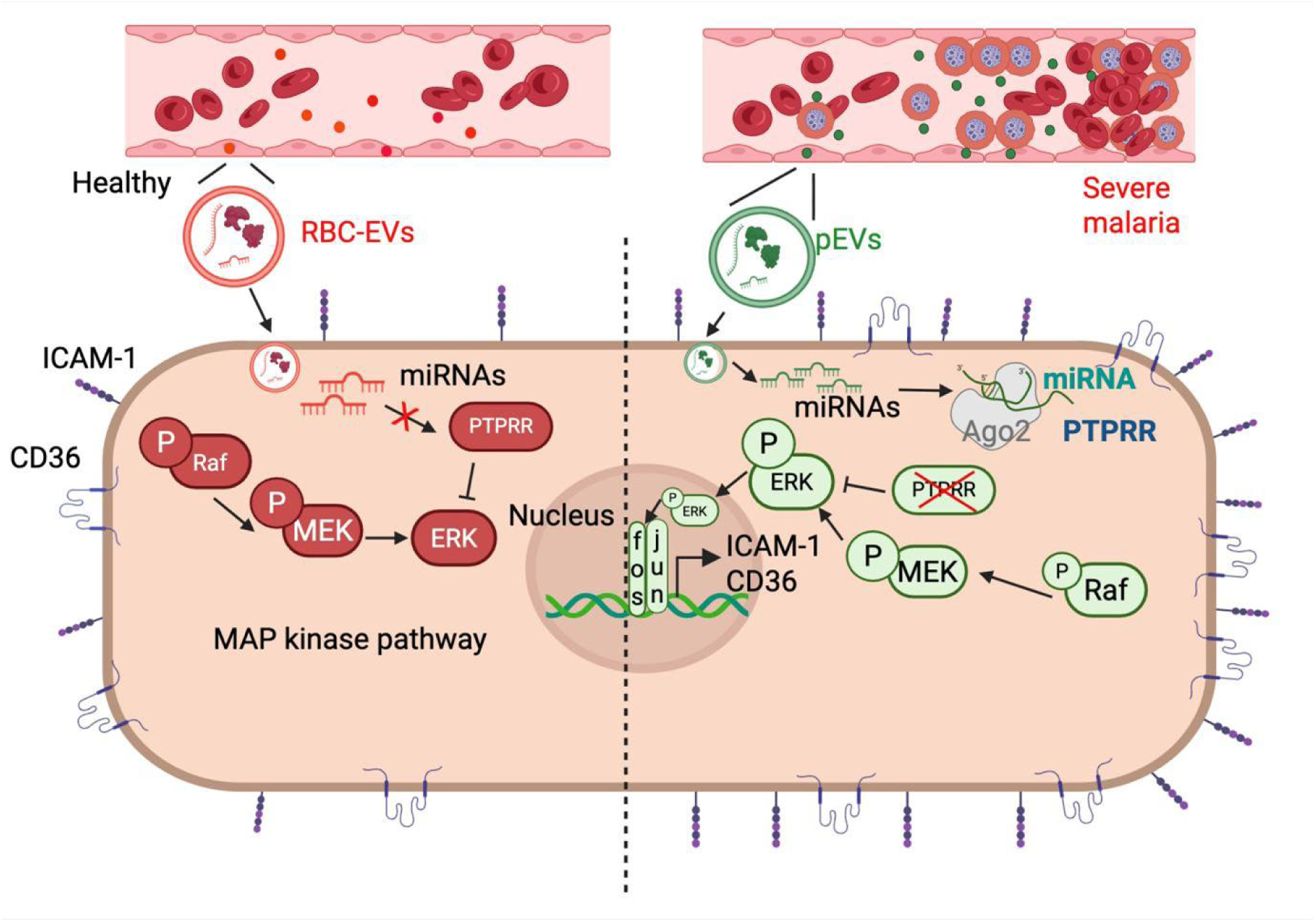
Model depicting the cross-talk between parasite and host through pEVs. pEVs upon uptake by host cells release miRNAs that target PTPRR, a phosphatase. The inhibition of PTPRR results in the accumulation of p-ERK that activates transcription of ICAM-1 and CD36 through c-fos and c-jun.

## Acknowledgments

MS is supported by UGC fellowship and SG is supported by DST-ECR/2017/001703, ANRF-SPG/2021/003333, DBT (BT/PR40722/MED/29/1529/2020) and IISER Tirupati core funding. Work in the SCB unit at OIST is funded by core subsidy from Okinawa Institute of Science and Technology Graduate University.

## Authors Contribution

S.G. and R.R.A. conceived study. M.S., R.R.A., S.G. conducted experiments. M.S., R.R.A. and S.G. analyzed data and discussed results. M.S., R.R.A. and S.G. wrote, read and approved the manuscript.

## References

[1] G.M.P. (gmp), World malaria report 2024, (2024), https://www.who.int/publications/i/item/9789240104440 (accessed July 10, 2025).

[2] M. Wahlgren, S. Goel, R. Akhouri, Variant surface antigens of Plasmodium falciparum and their roles in severe malaria, Nat. Rev Microbiol (2017); 15:479–491. 10.1038/nrmicro.2017.47.

[3] L.H. Miller, D.I. Baruch, K. Marsh, O.K. Doumbo. The pathogenic basis of malaria, Nature (2002); 415:673–679. 10.1038/415673a.

[4] B. Deb, A. Das, R. Vilvadrinath, A. Jangra, M.S. Shukla, R.R. Akhouri, et al. Glycophorin B-PfEMP1 interaction mediates robust rosetting in Plasmodium falciparum, Int. J. Biol. Macromol (2024); 262:129868. 10.1016/j.ijbiomac.2024.129868.

[5] A. Mahamar, O. Attaher, B. Swihart, A. Barry, B.S. Diarra, M.B. Kanoute, et al. Host factors that modify Plasmodium falciparum adhesion to endothelial receptors, Sci. Rep (2017); 7:13872. 10.1038/s41598-017-14351-7.

[6] L. Dos Santos Ortolan, M. K. Sercundes, G. C. Moura, T. de Castro Quirino, D. Debone, D. de Sousa Costa, et al. Endothelial protein C receptor could contribute to experimental malaria-associated acute respiratory distress syndrome, J. Immunol. Res. (2019); 2019:3105817. 10.1155/2019/3105817.

[7] A. Mayor, A. Hafiz, Q. Bassat, E. Rovira-Vallbona, S. Sanz, S. Machevo, et al. Association of severe malaria outcomes with platelet-mediated clumping and adhesion to a novel host receptor, PLoS One (2011); 6:e19422. 10.1371/journal.pone.0019422.

[8] J. Porta, A. Carota, G.P. Pizzolato, E. Wildi, M.C. Widmer, C. Margairaz, et al. Immunopathological changes in human cerebral malaria, Clin. Neuropathol (1993); 12:142–146. https://pubmed.ncbi.nlm.nih.gov/8100753/.

[9] A.K. Tripathi, D.J. Sullivan, M.F. Stins. Plasmodium falciparum-infected erythrocytes increase intercellular adhesion molecule 1 expression on brain endothelium through NF-kappaB, Infect. Immun (2006); 74:3262–3270. 10.1128/IAI.01625-05.

[10] J. Allweier, M. Bartels, H. Torabi, M.D.P.M. Tauler, N.G. Metwally, T. Roeder, et al. Cytoadhesion of Plasmodium falciparum-infected red blood cells changes the expression of cytokine-, histone- and antiviral protein-encoding genes in brain endothelial cells, Mol. Microbiol (2024); 122:948–967. 10.1111/mmi.15331.

[11] N. Regev-Rudzki, D.W. Wilson, T.G. Carvalho, X. Sisquella, B.M. Coleman, M. Rug, et al. Cell-cell communication between malaria-infected red blood cells via exosome-like vesicles, Cell (2013); 153 1120–1133. 10.1016/j.cell.2013.04.029.

[12] P.-Y. Mantel, D. Hjelmqvist, M. Walch, S. Kharoubi-Hess, S. Nilsson, D. Ravel, et al. Infected erythrocyte-derived extracellular vesicles alter vascular function via regulatory Ago2-miRNA complexes in malaria, Nat. Commun (2016); 7 (2016) 12727. 10.1038/ncomms12727.

[13] A. Abdi, L. Yu, D. Goulding, M.K. Rono, P. Bejon, J. Choudhary, et al. Proteomic analysis of extracellular vesicles from a Plasmodium falciparum Kenyan clinical isolate defines a core parasite secretome, Wellcome Open Res (2017); 2:50. 10.12688/wellcomeopenres.11910.2.

[14] P.-Y. Mantel, A.N. Hoang, I. Goldowitz, D. Potashnikova, B. Hamza, I. Vorobjev, et al. Malaria-infected erythrocyte-derived microvesicles mediate cellular communication within the parasite population and with the host immune system, Cell Host Microbe (2013);13 521–534. 10.1016/j.chom.2013.04.009.

[15] K.A. Babatunde, S. Mbagwu, M.A. Hernández-Castañeda, S.R. Adapa, M. Walch, L. Filgueira, et al. Malaria infected red blood cells release small regulatory RNAs through extracellular vesicles, Sci. Rep (2018). 8:884. 10.1038/s41598-018-19149-9.

[16] R.R. Akhouri, S. Goel, H. Furusho, U. Skoglund, M. Wahlgren. Architecture of human IgM in complex with P. falciparum erythrocyte membrane protein 1, Cell Reports (2016); 14:723–736. 10.1016/j.celrep.2015.12.067.

[17] Q. Guo, Y. Jin, X. Chen, X. Ye, X. Shen, M. Lin, et al. NF-κB in biology and targeted therapy: new insights and translational implications, Signal Transduct. Target. Ther (2024); 9:53. 10.1038/s41392-024-01757-9.

[18] Y. Fan, C. Liu, X. Qin, Y. Wang, Y. Han, Y. Zhou. The role of ERK1/2 signaling pathway in Nef protein upregulation of the expression of the intercellular adhesion molecule 1 in endothelial cells, Angiology (2010); 61:669–678. 10.1177/0003319710364215.

[19] D.-Y. Liang, F. Liu, J.-X. Chen, X.-L. He, Y.-L. Zhou, B.-X. Ge, et al.Porphyromonas gingivalis infected macrophages upregulate CD36 expression via ERK/NF-κB pathway, Cell. Signal (2016); 28:1292–1303. 10.1016/j.cellsig.2016.05.017.

[20] C. Zhang, X. Luo, J. Chen, B. Zhou, M. Yang, R. Liu, et al. Osteoprotegerin promotes liver steatosis by targeting the ERK-PPAR-γ-CD36 pathway, Diabetes (2019); 68:1902–1914. 10.2337/db18-1055.

[21] J. Zhu, X. Wu, S. Goel, N.M. Gowda, S. Kumar, G. Krishnegowda, et al. MAPK-activated protein kinase 2 differentially regulates plasmodium falciparum glycosylphosphatidylinositol-induced production of tumor necrosis factor-{alpha} and interleukin-12 in macrophages, J. Biol. Chem (2009). 284:15750–15761. 10.1074/jbc.M901111200.

[22] L.O. Murphy, S. Smith, R.-H. Chen, D.C. Fingar, J. Blenis. Molecular interpretation of ERK signal duration by immediate early gene products, Nat. Cell Biol (2002). 4:556–564. 10.1038/ncb822.

[23] T.D. Halazonetis, K. Georgopoulos, M.E. Greenberg, P. Leder. c-Jun dimerizes with itself and with c-Fos, forming complexes of different DNA binding affinities, Cell (1988);55:917–924. 10.1016/0092-8674(88)90147-x.

[24] M.E. Bahar, H.J. Kim, D.R. Kim. Targeting the RAS/RAF/MAPK pathway for cancer therapy: from mechanism to clinical studies, Signal Transduct. Target. Ther (2023); 8:455. 10.1038/s41392-023-01705-z.

[25] A. Bulle, P. Liu, K. Seehra, S. Bansod, Y. Chen, K. Zahra, et al. Combined KRAS-MAPK pathway inhibitors and HER2-directed drug conjugate is efficacious in pancreatic cancer, Nat. Commun (2024); 15:2503. 10.1038/s41467-024-46811-w.

[26] M.K. Singh, S. Altameemi, M. Lares, M.A. Newton, V. Setaluri. Role of dual specificity phosphatases (DUSPs) in melanoma cellular plasticity and drug resistance, Sci. Rep (2022); 12:14395. 10.1038/s41598-022-18578-x.

[27] S. Niture, B.H.M. Mooers, D.H. Wu, M. Hart, J. Jaboin, D. Seneviratne. Dual-specificity protein phosphatase 1: A potential therapeutic target in cancer, iScience (2025); 28:113706. 10.1016/j.isci.2025.113706.

[28] Z. Yao, R. Seger. The molecular mechanism of MAPK / ERK inactivation, Curr. Genomics(2004); 5:385–393, 10.2174/1389202043349309.

[29] T. Du, X. Hu, Z. Hou, W. Wang, S. You, M. Wang, et al. Re-expression of epigenetically silenced PTPRR by histone acetylation sensitizes RAS-mutant lung adenocarcinoma to SHP2 inhibition, Cell. Mol Life Sci (2024); 81:64. 10.1007/s00018-023-05034-w.

[30] L. Piatti, A. Batzilla, F. Nakaki, H. Fleckenstein, F. Korbmacher, R.K.M. Long, et al. Plasmodium falciparum egress disrupts endothelial junctions and activates JAK-STAT signaling in a microvascular 3D blood-brain barrier model, Nat. Commun (2025); 16:7262. 10.1038/s41467-025-62514-2.

[31] B. Othman, L. Zeef, T. Szestak, Z. Rchiad, J. Storm, C. Askonas, et al.Different PfEMP1-expressing Plasmodium falciparum variants induce divergent endothelial transcriptional responses during co-culture, PLoS One (2023);18:e0295053. 10.1371/journal.pone.0295053.

[32] D. Nantakomol, A.M. Dondorp, S. Krudsood, R. Udomsangpetch, K. Pattanapanyasat, V. Combes, et al. Circulating red cell-derived microparticles in human malaria, J. Infect. Dis (2011); 203:700–706. 10.1093/infdis/jiq104.

[33] Y. Wu, S. Leyk, H. Torabi, K. Höhn, B. Honecker, M.D.P.M. Tauler, et al. Plasmodium falciparum infection reshapes the human microRNA profiles of red blood cells and their extracellular vesicles, iScience (2023); 26:107119. 10.1016/j.isci.2023.107119.

[34] J.A. Kochanowsky, P.M. Mira, S. Elikaee, K. Muratore, A.K. Rai, A.M. Riestra, et al. Trichomonas vaginalis extracellular vesicles up-regulate and directly transfer adherence factors promoting host cell colonization, Proc. Natl. Acad. Sci. U. S. A (2024); 121:e2401159121. 10.1073/pnas.2401159121.

[35] G. Hatzivassiliou, J.R. Haling, H. Chen, K. Song, S. Price, R. Heald, et al.Mechanism of MEK inhibition determines efficacy in mutant KRAS- versus BRAF-driven cancers, Nature (2013); 501:232–236. 10.1038/nature12441.

[36] C. Ambrogio, J. Köhler, Z.-W. Zhou, H. Wang, R. Paranal, J. Li, et al. KRAS dimerization impacts MEK inhibitor sensitivity and oncogenic activity of mutant KRAS, Cell (2018); 172:857–868.e15. 10.1016/j.cell.2017.12.020.

[37] M. Cargnello, P.P. Roux. Activation and function of the MAPKs and their substrates, the MAPK-activated protein kinases, Microbiol. Mol. Biol. Rev (2011); 75:50–83. 10.1128/MMBR.00031-10.

[38] R. Lang, F.A.M. Raffi. Dual-specificity phosphatases in immunity and infection: An update, Int. J. Mol. Sci (2019); 20:2710. 10.3390/ijms20112710.

[39] J. Munkley, N.P. Lafferty, G. Kalna, C.N. Robson, H.Y. Leung, P. Rajan, et al. Androgen-regulation of the protein tyrosine phosphatase PTPRR activates ERK1/2 signalling in prostate cancer cells, BMC Cancer(2015); 15:9, 10.1186/s12885-015-1012-8.

